# Sorting of persistent morphological polymorphisms links paleobiological pattern to population process

**DOI:** 10.1101/2023.04.10.536283

**Authors:** Charles Tomomi Parins-Fukuchi

## Abstract

Biological variation fuels evolutionary change. Across longer timescales, however, polymorphisms at both the genomic and phenotypic levels often persists longer than would be expected under standard population genetic models such as positive selection or genetic drift. Explaining the maintenance of this variation within populations across long timespans via balancing selection has been a major triumph of theoretical population genetics and ecology. Although persistent polymorphisms can often be traced in fossil lineages over long periods through the rock record, paleobiology has had little to say about either the long-term maintenance of phenotypic variation or its macroevolutionary consequences. I explore the dynamics that occur when persistent polymorphisms maintained over long lineage durations are filtered into descendant lineages during periods of demographic upheaval that occur at speciation. I evaluate these patterns in two lineages: *Ectocion*, a genus of Eocene mammals, and botryocrinids, a Mississippian cladid crinoid family. Following origination, descendants are less variable than their ancestors. The patterns by which ancestral variation is sorted cannot be distinguished from drift. Maintained and accumulated polymorphisms in highly variable ancestral lineages such as *Barycrinus rhombiferus* (Owen and Shumard 1852) may fuel radiations as character states are sorted into multiple descendant lineages. Interrogating the conditions under which trans-specific polymorphism is either maintained or lost during periods of demographic and ecological upheaval can explain how population-level processes contribute to the emergent macroevolutionary dynamics that shape the history of life as preserved in the fossil record.

**Non-technical abstract:** Understanding how morphological variation changes within populations over relatively short timescales in response to environmental changes and ecology (i.e., thousands of years) is a major focus of paleontology and evolutionary biology. A distinct focus is in understanding the broad-scale patterns by which lineages have diversified into distinct environments over geologic time (i.e., millions of years). One major challenge has been reconciling how and whether processes acting over shorter timescales shape the patterns observed over long timescales. One way of examining morphological variation at the population level is by examining the distribution of polymorphic character states--discrete anatomical features that vary within a population. Fossil species often maintain such polymorphisms for long periods of time, with such variation even sometimes inherited by new species from their ancestors. In this article, I suggest that examining how these polymorphisms are distributed among incipient descendant lineages might help link the ecological and evolutionary processes that act at the population level (e.g., natural selection, genetic drift, competition, predation) to the paleobiological patterns that are often reconstructed across many species and over long timescales. I explore these dynamics in two lineages: *Ectocion*, a genus of Eocene mammals, and botryocrinids, a Mississippian cladid crinoid family. I found that new lineages typically have fewer polymorphisms than their ancestors, suggesting that ancestral variation is “sorted” into incipient lineages during speciation. This variation appears to be sorted randomly, which means that it is not possible to detect the influence of natural selection in guiding the inheritance of ancestral morphologies. I suggest that the patterns by which ancestral variation is sorted into new species may explain patterns of lineage diversification over long timescales, highlighting how population processes can extend their influence over longer timescales to shape large-scale evolutionary dynamics.

## Introduction

Polymorphism is ubiquitous within natural populations. Spanning levels from the genome to the phenotype, it is difficult to look far without observing variation. Positive selection, purifying selection, and genetic drift are all expected to remove variation from the population, which has appeared at odds with the vast reservoirs of variation displayed by many organisms. However, theoretical and empirical work has shown how environmental heterogeneity (McDonald and Ayala 1974, Christiansen 1975, Chakraborty and Fry 2016, Gallet et al. 2018) and negative frequency-dependent selection (NFDS) (Clark 1979, Charlesworth 2006, Fitzpatrick et al. 2007, Takahashi et al. 2010) can protect such “balanced polymorphisms”, maintaining genetic and phenotypic variation over long timescales. While variation within populations is most often examined from the standpoint of microevolution, there is increasing evidence that polymorphisms can persist over long time periods. By spanning temporal and taxonomic scales, persistent polymorphic variation might offer a window into understanding how the microevolutionary processes that promote long term polymorphism might shape macroevolutionary patterns and processes.

Despite its omnipresence in nature and frequent treatment in population genetics, the causes and effects of intra-population polymorphism have been understudied at deeper timescales. Characters employed in phylogenetic studies frequently display polymorphism across multiple lineages but have frequently downplayed this variation. Nevertheless, they are sufficiently common to substantially impact phylogenetic inferences when not explicitly accommodated in analyses (Wiens 1995, 1999). Phenotypic polymorphisms also feature strongly in paleontological systematics. Characters chosen for cladistic analysis between fossil lineages are frequently rife with polymorphism, which can present challenges to existing algorithms (e.g., Trueman 2010, Gilbert 2013, Whiting et al., 2016, plus many others). Relative to the goals of systematics, these are often treated as nuisances. This is perhaps due to a historical gaps in the integration between systematics, phylogenetic methods, and population genetics. Despite being frequently observed, the apparent maintenance of polymorphism across populations separated by millions of years of independent evolution remains understudied as a macroevolutionary phenomenon.

A small, but growing, body of work has contributed early inklings of the potential for sustained variation to explain evolutionary dynamics across species boundaries and over increasingly deep timescales. One increasingly important issue in evolutionary ecology concerns how phenotypic and genetic variation evolves across species boundaries (e.g., Thompson et al. 2019). Standing genetic variation can facilitate rapid adaptation when populations are exposed to new ecological pressures, such as might occur during ecological speciation (Barrett and Schluter 2008, Sicard et al. 2016, Lai et al. 2019, McGee et al. 2020). There has also been an increasing focus in population genomics upon understanding how (often balanced) polymorphisms in large ancestral populations may be “sieved” during the speciation process, becoming fixed among descendants either randomly or by selection (Pease et al. 2016, Guerrero and Hahn 2017). “Deeply persistent” polymorphisms can even be maintained by NFDS over millions of years, shaping macroevolutionary dynamics over large clades (Igic et al. 2006, Goldberg et al. 2010).

Paleobiologists have spent less time seeking process-based explanations for the evolution of phenotypic variation. However, the fossil record might be leveraged to offer unique insight into this fundamental question. Fossilized populations often display maintain variation over long intervals of time (e.g., Brothwell 1963, also see examples reviewed extensively by Van Valen 1969). Although the fossil record has been underutilized as a laboratory to investigate population genetics, the marine invertebrate record has shown promise in its ability to illuminate the processes governing the maintenance of phenotypic variation (Kermack 1954). Intraspecific variation has been hypothesized to have contributed to the explosive diversification of trilobites during the Cambrian (Webster 2007). Fossil lineages frequently display high population-level variation, both in quantitative traits and discrete character states. This variation can be maintained for long periods of time, across a range of environmental conditions (including environmental stasis—see Schopf and Gooch 1972). Despite these substantial advances, the full incorporation of persistent, trans-specific phenotypic polymorphisms in the fossil record into paleobiological theory has remained incomplete. Nevertheless, the presence of persistent intraspecific variation shared between fossilized lineages is an often-unrecognized curiosity when viewed from the lens of population genetics that may contain hidden insights when bridging micro- and macroevolutionary scales.

While not always formally recognized as such, the evolution of phenotypic variation during speciation has long factored implicitly into fundamental questions in paleobiology (Simpson 1944, Eldredge and Gould 1972, Gingerich 1974, Polly 1997). In particular, paleobiologists have often been concerned with understanding how speciation proceeds between ancestral and descendant lineages. Key neontological work has also suggested that examining the maintenance of ancestral variation and its distribution among descendant populations may explain the formation and adaptive divergence of new lineages (Wright 1982, Schluter and Conte 2009). Budding speciation, identified as asymmetric speciation between a large ancestral population, and a smaller descendant population, has long served as paleobiological bread and butter (e.g., Rensch 1959, Jackson and Cheetham 1994, Aze et al. 2011, Warnock et al. 2020, Parins-Fukuchi 2021). One important characteristic of budding is the persistence of the ancestral lineage alongside its newly formed descendant. There may be a geographic component, wherein the ancestral lineage occupies a wide geographic range and the descendant lineage arises due to peripheral isolation within a smaller area, but this is not essential. Budding may also be used to describe the speciation dynamics observed in important neontological work (e.g., Schluter 2000), and so is a possible conceptual bridge by which the population processes underlying the evolution of variation studied by neontologists may be more thoroughly integrated into the paleobiological research program.

Tracing the evolution of polymorphic characters across the repeated episodes of budding speciation found in dense fossil records would provide a unique opportunity to test the predictions from population genetics surrounding the inheritance of persistent, standing variation into descendant populations and examine their effects over longer time scales. When populations bud from an ancestral lineage, the pattern by which traits and alleles become fixed in the new species may be “parallel”, “divergent”, or “random” (Schluter and Nagel 1995, Thompson 2019). In the parallel case, new descendant lineages tend to fix the same traits as they radiate into similar environments. In the divergent case, descendants evolve in opposing phenotypic directions as they radiate into distinct environments. Lastly, newly budded species may randomly sort variation present in the ancestor if demographic asymmetry in population divergences leads to bottlenecks. Each of these leads to different implications both within lineages and over deeper time.

In this paper, I attempt to explain the evolution of phenotypic variation across episodes of budding speciation in two fossil lineages: *Ectocion*, a genus of Eocene mammals closely related to perissodactyla; and botryocrinids, a family of mostly Mississippian cladid crinoids. My first goal was to explain how an explicit treatment of persistent polymorphisms can clarify interpretations of biological patterns in the fossil record. I then sought to further examine the ways by which persistent ancestral variation is sorted into descendant lineages. I sought to test whether 1) descendant species tended to be less variable than ancestral species, a key prediction of budding speciation, and 2) whether ancestral variation tended to sort randomly, consistent with the fixation of polymorphisms due to population bottlenecking, or non-randomly, consistent with either convergent or divergent selection following speciation.

## Materials and Methods

### Data and code availability

All data and code used in the analyses presented here are available on Figshare and associated with the digital object identifier 10.6084/m9.figshare.22584097.

### Datasets

I harvested morphologic and stratigraphic datasets encompassing two lineages from the literature: Ectocion (Thewissen 1992) and botryocrinidae (Gahn and Kammer 2002). These two lineages offer several advantages. Both possess dense fossil records, simplifying the identification of likely ancestor-descendant relationships. They have also benefited from careful work delimiting species and tracing continuous lineages through stratigraphic zones in particularly well-studied regional faunas. The OTUs represented within each dataset can therefore likely be regarded as trustworthy. This careful work has also provided greater certainty in trusting the persistence of polymorphisms observed in each dataset—while character state frequencies among several polymorphic characters fluctuates throughout the range of each lineage, overall, the polymorphic variation attributed to each lineage largely remained intact. The maintenance of polymorphism throughout lineage durations is further supported by the persistence of polymorphic characters across several speciations. The repeated observation of polymorphism across multiple lineages favors the persistence of polymorphisms within and between lineages as the most parsimonious explanation.

The *Ectocion* dataset was entirely dental. After reconstructing the phylogeny, I mapped the evolution of the P3 metacone, which was polymorphic across two lineages over long time spans. While explicitly mapping such detailed dental morphology to dietary function is a very challenging prospect, it is reasonable to imagine that the metacone, which is haphazardly distributed across *Ectocion*, may have provided some dimension of dietary function. The botryocrinid dataset sampled characters across the bauplan, spanning several anatomical regions, including the arms, stalk, and calyx plating, including configuration of the posterior plates. Polymorphisms were observed across all regions. Functional interpretations for each of these traits is very challenging. However, all, or nearly all, of them do interact with the environment in some way. Premolar morphology undoubtedly plays a major role in food processing in terrestrial vertebrates, such as *Ectocion*. Functional morphology is also well characterized in crinoids. For example, crown morphology appears to contribute to hydrodynamics (Cole et al. 2019), while column morphology can correspond to the ecological niche of crinoid species via tiering (Ausich 1980, Ausich and Bottjer 1982). As a result, it is feasible that they could be under some form of selection, although closer biomechanical and eco-phenotypic analysis would be needed to better understand the specific functional context of each of these traits within the lineages examined.

Several of the characters observed by Gahn and Kammer (2002) were not true discrete characters, but rather, discretized descriptions of variation that is fundamentally continuous. While this challenges interpretations of these particular characters within the framework of strict allelic polymorphism, I believe they were appropriate to retain in the current study. This is because my goal in analyzing the botyrocrinid dataset was in examining how ancestral variation is sorted and maintained across ancestor-descendant transitions. While coarse, even these discretized treatments offer resolution into this topic, especially if the continuous variation represented within is at least somewhat discontinuous in its frequency distribution. Such variation is also likely to correspond to higher heterozygosity among underlying loci, supporting their use in interpreting broad patterns underlying the evolution of variation.

### Phylogenetic reconstruction

I implemented a very simple method for the reconstruction of phylogeny, including ancestor-descendant relationships, for this study. It is based upon the greedy, agglomorative algorithm employed by neighbor-joining (Saitou and Nei 1987). It only differs by incorporating stratigraphic information into the distance matrix and allowing earlier-occurring lineages to serve as direct ancestors of those that occur later. The approach is also very similar to the greedy algorithm described by Alroy (1995), with some simplifications. It starts by constructing a matrix of all of the morphological distances between each OTU, calculated as the number of distinct character states between each possible OTU pair. In the case of polymorphism, OTU pairs are assigned a distance of zero if the intersection between character states displayed by each was non-empty. Stratigraphic distances are then added. These are calculated as the number of discrete gaps separating non-overlapping lineages. Lineages with overlapping or abutting ranges are assigned a stratigraphic distance of zero. The least dissimilar OTU pair is then identified and joined. If one of the OTUs first occurs before the other, it is assumed to be ancestor. If they start in the same time horizon, they are assigned a shared hypothetical ancestor that is placed in the same time bin. The algorithm then proceeds iteratively, adding OTU pairs until all pairs are joined. The tree is then rooted using either an outgroup or by presuming the most stratigraphically basal OTU is the root. The resulting tree is one that minimizes the amount of evolutionary change and stratigraphic gaps across lineages. It thus bears similarities to both stratophenetics (Gingerich 1979) and stratocladistics (Fisher 2008).

The method used here is very simplistic. I do not endorse its use in large, complex datasets. Nevertheless, I believe it is adequate for my purposes here. This is because the datasets employed in this study are quite small and benefit from having very dense and well-characterized stratigraphic and geographic ranges. In addition, the high degree of polymorphism displayed by both lineages supported the use of the distance approach used here over existing implementations in that it offered a simple solution to the treatment of polymorphism when calculating evolutionary distances. This was important given the high degree of polymorphism in the *Barycrinus* dataset and the inability of most existing parametric and parsimony approaches to accommodate polymorphism explicitly. Several existing Bayesian approaches entertain ancestor-descendant hypotheses, including budding speciation (Stadler et al. 2018). However, given the extensive challenges associated with searching treespace to explicitly identify such hypotheses, rather than simply integrating over them, I felt the simplicity of the method here was justified for use on the particular datasets employed here. As a final note, in the absence of contrary information, it is more parsimonious to assume that taxa are related through ancestor-descendant sequences rather than invoking hypothetical ancestors (Polly 1997). This is particularly the case if budding speciation is assumed to predominate and sampling rates are high (Foote 1996).

Moderate to low preservation, when paired with bifurcating speciation, may make hypothetical ancestors a safer *a priori* assumption, and so the extent to which this appeal to model parsimony generalizes to other cases should be gauged against theses considerations. Nevertheless, given that the density of the fossil records displayed by the lineages examined here increases the odds of sampling ancestral taxa, this reasoning should lend some epistemological confidence in the method’s behavior of minimizing the number of hypothetical ancestors. Nevertheless, my approach to phylogeny reconstruction, while likely appropriate for the densely sampled and carefully studied lineages employed here, is propositional, and ultimately somewhat crude, in its nature. It should therefore be accompanied by a more rigorous, evaluative, approach for future larger-scale studies of budding dynamics in the fossil record, such as stratocladistics (e.g., Fisher 1991, 2008) stratolikelihood (e.g., Wagner 1998, 2000), or full-Bayesian approaches (e.g., Wright et al. 2021). On larger datasets, the simple approach implemented here may remain useful in generating a starting tree for more exhaustive heuristic searches.

I applied this algorithm to reconstruct relationships in the taxa *Ectocion* and botryocrinids. Stratigraphic ranges were harvested from the same references from which I sourced the morphologic information. One small adjustment was made in the botryocrinids. I allowed *Barycrinus rhombiferus* (Owen and Shumard 1852) to be ancestral to *Barycrinus spurius* (Hall 1858), despite them having originated in the same layer. This is because Gahn and Krammer (2002) made a compelling case for *B. rhombiferus’* likely status as an ancestor to multiple other *Barycrinus* lineages. In addition, while *B. rhombiferus* and *B. spurius* were both long-lasting and highly polymorphic lineages, *B. rhombiferus* possess fewer derived character states than *B. spurius*. This makes a *B. rhombiferus* -> *B. spurius* ancestor-descendant pair more parsimonious, according to stratocladistic criteria, than the reverse. In addition, allowing the possibility also achieved a result with several fewer hypothetical ancestors, making a strong appeal to model parsimony. It should be clarified that *B. rhombiferus* was not, *a priori*, constrained or assumed to be ancestral to *B. spurius*. This possibility was simply *allowed* by the analysis.

### Random vs. non-random sorting of ancestral variation

I devised a simple permutation test to identify whether ancestral variation in botryocrinids tended to sort randomly among descendant lineages, which would be suggestive of genetic drift driving fixation during speciation bottlenecks (i.e., founder events), or whether descendant lineages tended to fix the same ancestral variants, which would be consistent with natural selection during adaptation in similar environments (i.e., ‘ecological speciation’ *sensu* Schluter and Conte 2009). Using the ancestor-descendant transitions reconstructed across *Barycrinus*, I took the polymorphic character states displayed by each ancestral lineage. I then randomly sampled a single character state from each ancestrally-polymorphic character to generate a hypothetical character sequence of inherited polymorphisms for each descendant lineage observed in the dataset. I then compared the pairwise Hamming distances between the simulated character states displayed by each descendant. To calculate distances, polymorphisms were treated as a unique character state. As a result, a comparison between a (01) polymorphism and (1) monomorphism resulted in a distance of 1. This was done according to the reasoning that, if incipient species were experiencing positive selection in new environments, they should purge the same ancestral character states.

My expectation was that descendants adapting to similar environments would likely share more character states, and thus have fewer differences. I repeated this sampling procedure 1000 times to generate a frequency distribution of pairwise distances between the character states displayed by each descendant that were randomly sampled from the polymorphic ancestor. I then compared the observed distances to the empirical distribution to identify any descendant taxon pairs that were more similar (indicating parallel adaptation) or different (indicating divergent adaptation) than expected under random sorting, using the 2.5% quantile as a significance threshold.

## Results and Discussion

### Ectocion phylogeny

Phylogenetic relationships reconstructed within *Ectocion* (Fig. 1) are largely concordant with the interpretation that Thewissen (1992) derived from the same dataset based on stratocladistic criteria. The main point of contention lies in the placement of *Ectocion mediotuber* (Thewissen 1990) and *Ectocion cedrus* (Thewissen 1990) in relation to *Ectocion collinus* (Russell 1929). The original study suggested the three taxa to form a grade, separated by hypothetical ancestors, whereas the current reconstruction posits that *E. cedrus* and *E. mediotuber* independently budded from *E. collinus*. This is a minor difference that will be discussed further below. Otherwise, the ancestor-descendant linking of *E. mediotuber* to *E. osbornianus* (Cope 1882) and *E. parvus* (Granger 1915) corresponds identically to the original study. Thewissen was conservative in his interpretation of *E. superstes*. However, the approach here suggested that it may have speciated from *E. osbornianus*. It should be noted that, while the diagram presented here gives the appearance of a direct ancestor-descendant relationship between the two, it is very possible that their true genealogical relationship is separated by one or more unobserved species. This would still make *E. osbornianus* an indirect ancestor.

**Figure 1.**
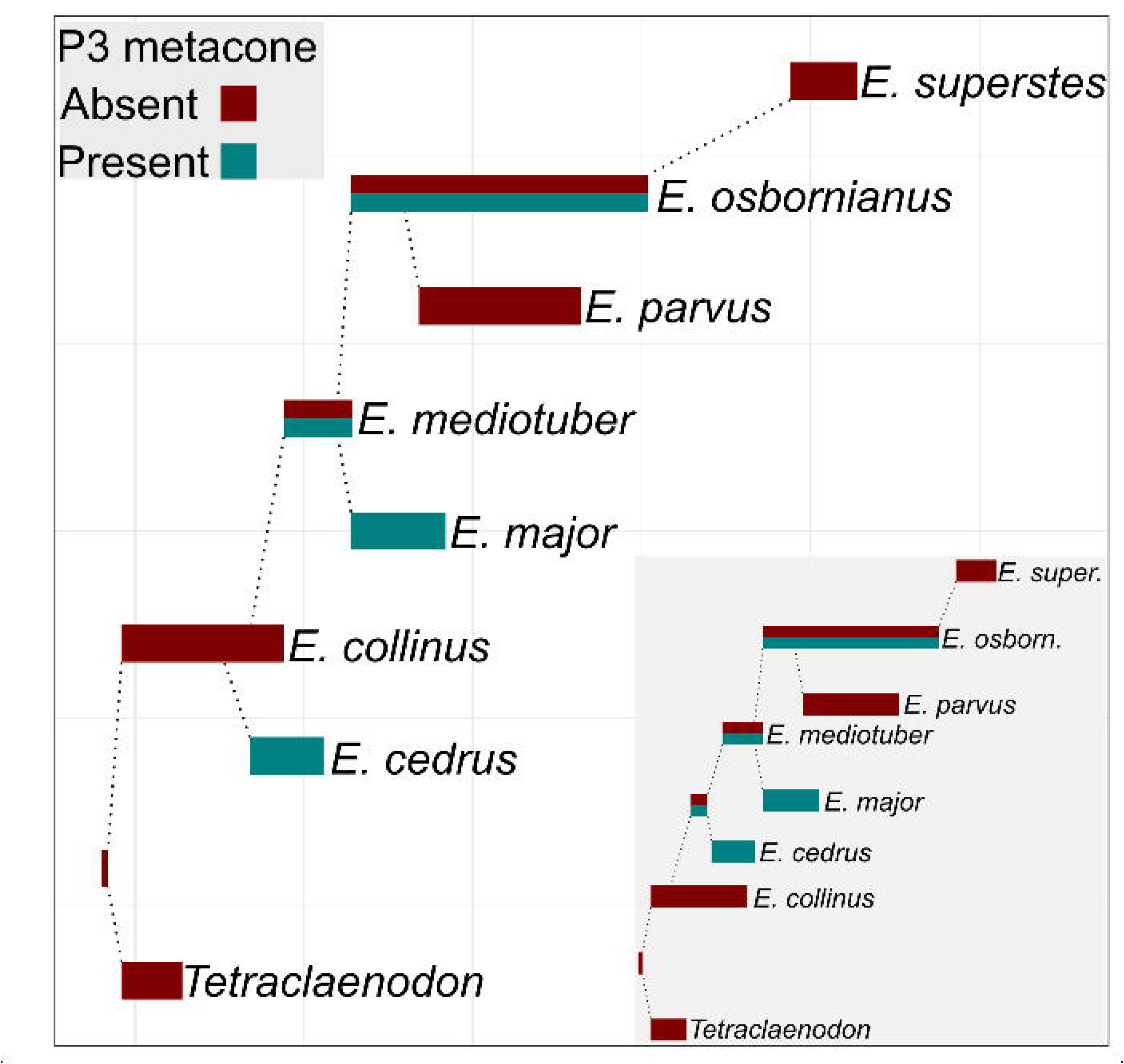
Reconstructed ancestor-descendant relationships within *Ectocion*. Bars represent stratigraphic ranges. Dotted lines indicate genealogical relationship between ancestral and descendant lineages. Colors reflect presence or absence of the metacone on the P3. Inset phylogeny represents hypothetical alternative reconstruction that better accommodates pattern in the evolution of P3 polymorphism. Further evaluation would be needed to distinguish between these two. Timescale reflects the discrete zonation used by Thewissen (1992).

### Evolution of the P3 metacone polymorphism

Mapping the evolution of the P3 metacone to the *Ectocion* phylogeny illustrates the importance of considering both ancestor-descendant relationships and polymorphism when reconstructing patterns in morphological evolution in the fossil record. A standard cladogram would fail to capture how polymorphisms in *E. mediotuber* and *E. osbornianus* persisted and sorted among the respective descendants of each, instead representing the pattern of character changes through multiple character reversals. One slightly confusing pattern in P3 metacone evolution is the apparent sudden emergence of a monomorphic metacone in *E. cedrus*. A more parsimonious interpretation, based on P3 morphology, of the sequence of lineage divergence than that achieved using the greedy method implemented here is given in the inset to Figure 1, which implies that the metacone evolved one time and sorted among descendant lineages while being maintained across multiple species. Alternatively, it is possible that the metacone truly did evolve twice in *Ectocion*. A stronger understanding of the developmental programs controlling dental variation would help to distinguish whether the metacone is sufficiently evolvable to have emerged twice, but distinguishing between these two possibilities falls outside the scope of this study. While the invocation of an additional, unaccounted polymorphic hypothetical ancestor provides a better explanation for P3 morphology evolution, detailed phylogenetic work that more thoroughly explores treespace using parsimony or probabilistic criteria will be needed to confidently distinguish between these possibilities.

Distinguishing between inheritance of ancestral polymorphism and character reversal is important for deriving meaningful biological interpretations. The pattern in P3 evolution reconstructed here is consistent with a scenario in which balancing selection maintains variation across a long-lived lineage, giving rise to descendant lineages that fix that ancestral variation in different ways. If the maintenance of polymorphism by balancing selection is as common as ecological theory might suggest, many character distributions among fossil taxa (which are often interpreted as being highly homoplaseous) may simply be driven by the sorting of variation among a small handful of variable ancestral lineages. In addition to potentially helping solve problems in paleontological systematics, exploration of these dynamics may shed light on the processes that drive the diversification of clades over paleobiological timescales.

The persistence of the P3 metacone polymorphism across *E. mediotuber* and *E. osbornianus* over several million years is difficult to explain by genetic drift alone. While some fluctuation was observed in the character state frequency throughout the range of *E. osbornianus*, the polymorphism remained at reasonably intermediate frequency, with >30% and <60% of individuals displaying the metacone throughout the duration of the lineage (Thewissen 1992). NFDS provides one possible explanation for this maintenance. Competition for food resources could provide negative, frequency-dependent feedback on the frequency of each P3 morph if metacone presence yielded different fitness outcomes related to the processing of some limiting food resource as a function of population density. Several modes of competition have long been recognized as an important cause of NFDS (Antonovics and Kareiva 1988, Dijikstra and Border 2018). The most relevant mode for P3 morphology might be resource competition associated with density-dependence. Alternatively, the persistence of the P3 polymorphism might instead result from some form of bet-hedging or response to soft selection in the presence of environmental heterogeneity over either space or time. At least equally likely is the possibility that P3 morphology is selectively neutral and fluctuated in frequency due to genetic drift. Whatever maintained this polymorphism, the results here demonstrate how sorting from ancestral stock shape the phenotypes displayed by incipient descendant lineages. The fixation of monomorphic P3 forms in the descendant lineages *E. major*, *E. parvus*, and *E. superstes* reflects either adaptation or drift from standing variation present in the ancestral lineages. If some form of balancing selection maintained P3 polymorphism in *E. mediotuber* and *E. osbornianus*, it is possible that demographic bottlenecks encountered during speciation of the three descendant lineages lead to random fixation. More exploration into the overall levels of phenotypic diversity and geography of these descendant lineages would help to examine the feasibility of this explanation.

### Barycrinus phylogeny

Throughout its evolutionary history, *Barycrinus* displayed several long-lived lineages that gave rise to more than one descendant (Fig. 2). In particular, *B. rhombiferus* is identified as having been a highly prolific lineage, with four immediate descendants. This is consistent with its status in Gahn and Kammer’s (2002) analysis, which found the placement of *B. rhombiferus* to be highly uncertain, leading the authors to identify the lineage as a “rogue taxon”. The authors interpreted this to suggest that this uncertainty may have resulted from *B. rhombiferus* having itself given rise to multiple later-occurring *Barycrinus* lineages. Based on patterns in the inheritance of ancestral character states, Gahn and Kammer suggested that *B. rhombiferus* was likely the direct ancestor of *B. magister* (Hall 1958), *B. spectabilis* (Meek and Worthen 1870), and *B. scitulus* (Meek and Worthen 1860)--an assertion that was further supported by Gahn (2003). They also suggested a possible close link between *B. rhombiferus* and *B. spurius*, going on to speculate that *B. rhombiferus* may be ancestral to most other *Barycrinus* lineages. Nevertheless, Gahn and Kammer’s analysis, which was relied only on traditional cladistic methods, achieved only low resolvion, perhaps due both to the rampant polymorphism and the complex pattern of ancestor-descendant relationships found in the clade. The phylogenetic tree recovered here is also largely consistent with quantitative results achieved using stratocladistics (Gahn 2003, pers. comm.) and Bayesian methods (Wright 2022, pers. comm.).

**Figure 2.**
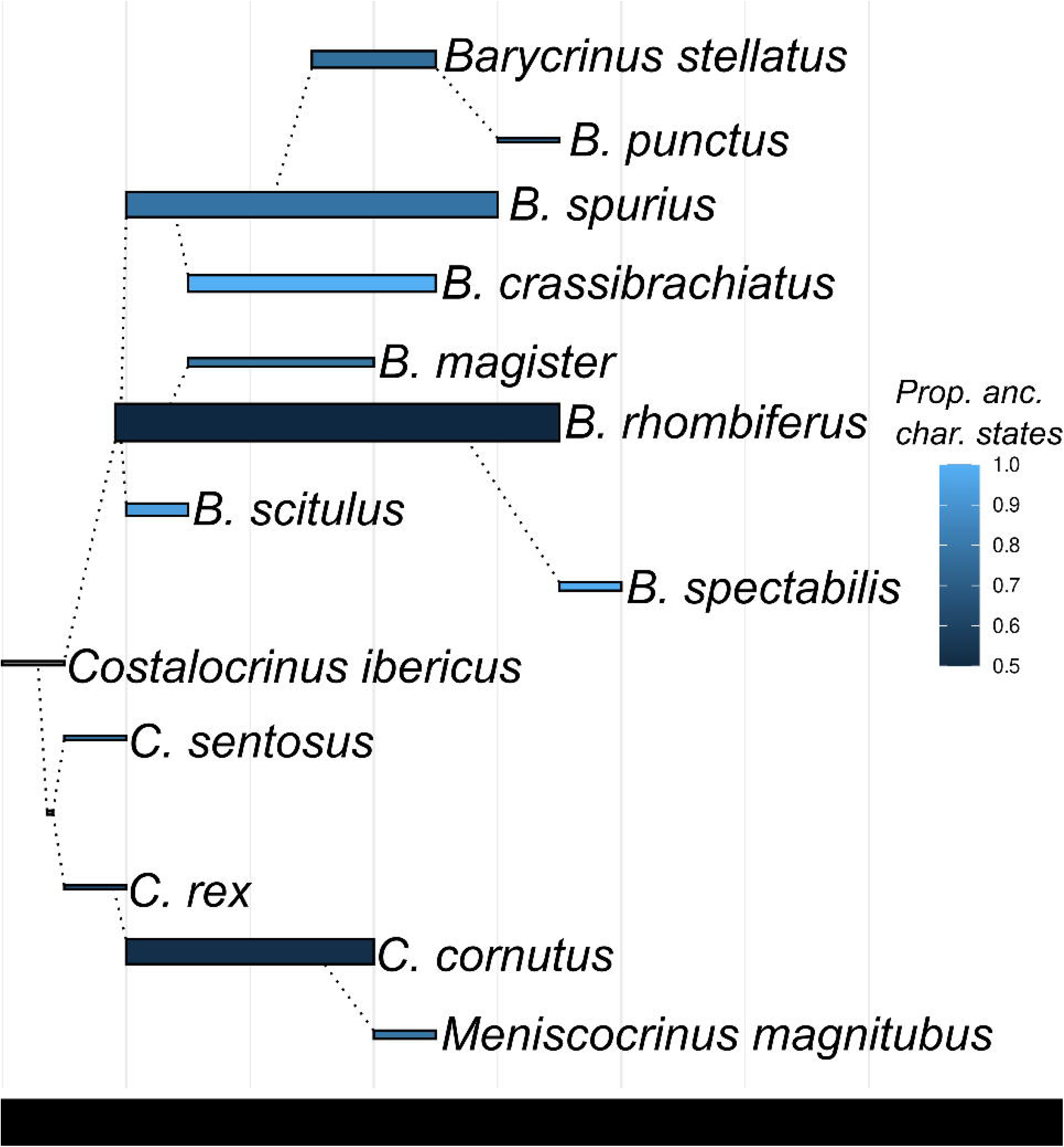
Phylogeny of Barycrinus. Bars represent stratigraphic ranges. Dotted lines indicate genealogical relationship between ancestral and descendant lineages. Width of stratigraphic lines represents scaled number of polymorphisms—a measure of genetic variation within each lineage. Shading represents the proportion of character states displayed by each lineage that were also possessed by its ancestor. Timescale approximates the discrete stratigraphic units used by Gahn and Kammer (2002).

*Barycrinus* underwent a rapid radiation while drawing upon a stock of ancestral variation maintained by *B. rhombiferus* and, to a lesser extent, *B. spurius*. This dynamic provides a small-scale demonstration of how the filtering of phenotypic variation during speciation might shape patterns at deeper timescales. In the phylogenetic analysis presented here, *B. rhombiferus* is depicted as having given rise to four descendant lineages. Two of these, *B. spectabilis* and *B. scitulus*, displayed a small fraction of the variation displayed by their ancestor as well as significantly reduced durations. If each of these incipient lineages occupied the same ecological landscape as *B. rhombiferus*, they may have been at a competitive disadvantage (at the lineage level) which would explain their shorter durations. The high phenotypic variability displayed by *B. rhombiferus* could stem from environmental heterogeneity caused by broader niche occupancy. If so, the increased survivorship displayed by generalist crinoids (Kammer et al. 1997, Cole 2021) might be explained by increased bet-hedging capability displayed by generalists that are able to maintain pools of phenotypic variation. Alternatively, if functional interpretations for the traits that are polymorphic in *B. rhombiferus* are more consistent with non-adaptive explanations (as may be the case--Gahn, personal communication), it is possible that its high variability simply stems from its increased ability to acquire (and subsequently maintain) variation over its long duration (Foote, personal communication). This explanation would require a scenario wherein mutation and genetic drift cooperate to introduce and maintain neutral variation over geologic timescales.

*B. magister* and *B. spurius* experienced longer durations than their siblings. This could be explained in *B. magister* by its evolution of additional derived character states not present in *B. rhombiferus*, which may be indicative of an escape to distinct ecological conditions. However, this scenario cannot be evaluated at present due to the geographic co-occurrence of both lineages and the lack of complete stem preservation in *B. magister* (Gahn, personal communication). While crinoids, as generalist suspension feeders, often occupy very similar ecological niches, a spectrum of niche differentiation does exist (Ausich 1980, Ausich and Bottjer 1982). The extended duration and multiple speciations produced by *B. spurius* might be explained by its continued maintenance of much of the variation possessed by *B. rhombiferus* as well as its derivation of several new characters. These possibilities remain speculative within the context of this article, however, they may be fruitfully explored as hypotheses to be tested in future work that includes more comprehensive ecological information that can be compared against functional morphological reconstructions. Evaluating the ideas presented here in light of such additional information may help to more rigorously explain the distinct causes underlying the patterns of lineage persistence and morphological variation displayed by *Barycrinus*. The analysis here represent only a starting point from which to draw deeper links between the evolution of phenotypic variation across species bounds and the ecological strategies employed by crinoid species.

### Random versus adaptive sorting of ancestral variation

The process of budding speciation, as typically conceived, may often yield population bottlenecks. If the bottleneck is strong enough, different character states would be expected to fix when small sub-populations are drawn from polymorphic ancestral populations over repeated trials (Fig. 3a). Alternatively, sufficiently strong positive selection would fix the same character state over repeated trials if incipient lineages repeatedly move out of the ancestral niche and into the same derived niche (Fig. 3b). I leveraged the repeated budding speciations displayed across *Barycrinus* to identify whether ancestral variation tended to sort randomly (Fig. 3a) or adaptively (Fig. 3b). Based on the permutation tests, it was not possible to distinguish the pattern by which the descendants of *B. rhombiferus* and *B. spurius*, sorted and fixed variation from their respective ancestors from the expectation under drift induced by bottlenecking (Tables 1 and 2). None of the pairs of descendants of either lineage showed statistically significant signs of parallel nor divergent speciation. Nevertheless, *B. spectabilis* and *B. magister* were somewhat more similar than expected under random sorting, while *B. spectabilis* and *B. spurius* were more divergent than expected. The lack of significance could be a consequence of the small pool of traits included in this dataset and so further tests are needed to see whether more comprehensive phenotypic sampling would distinguish the observed patterns from drift.

**Figure 3.**
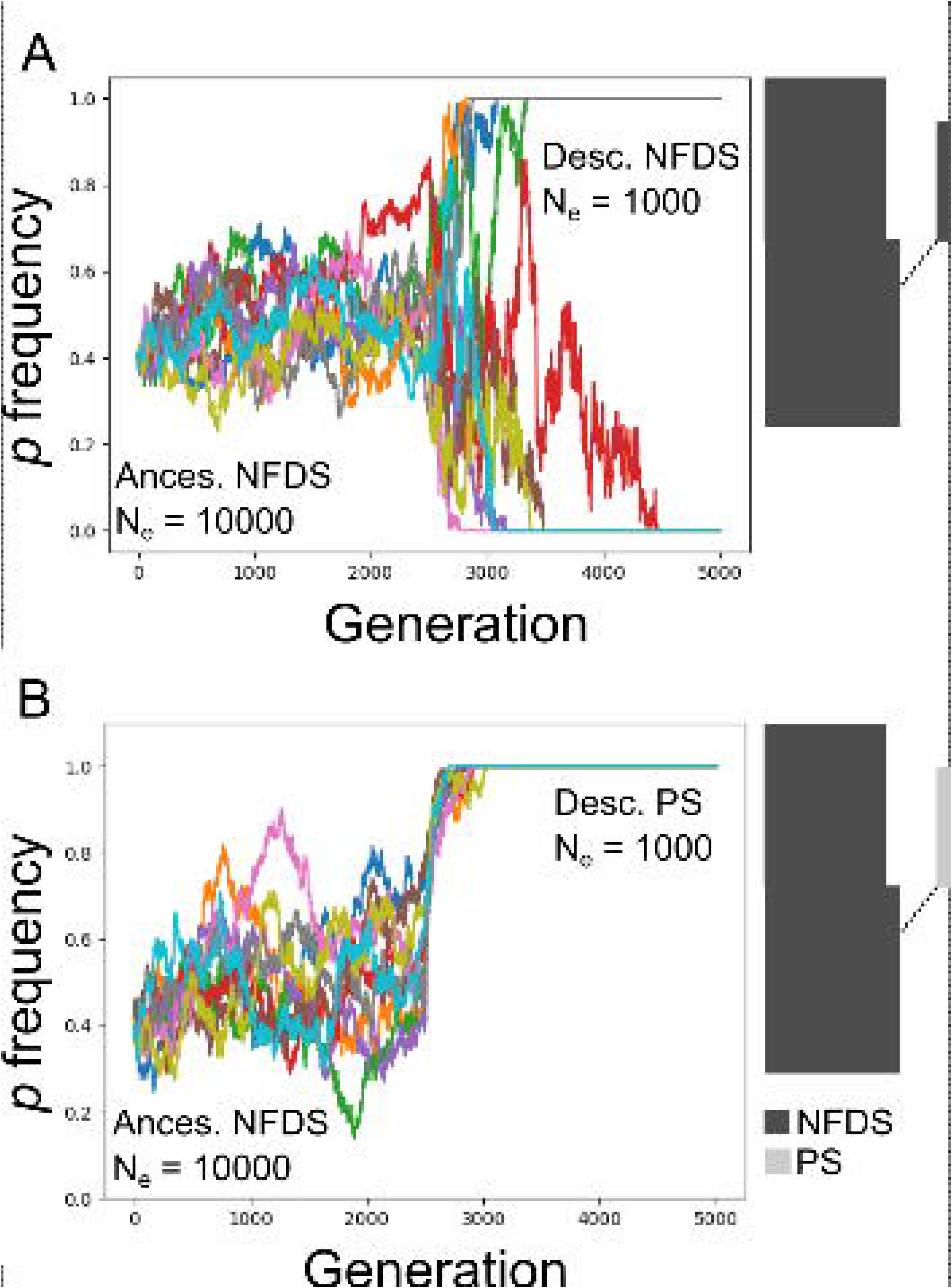
Distribution of simulated allele frequencies over 10 replicated populations while a balanced ancestral polymorphism is filtered into a budding descendant under two different adaptive scenarios: A) Polymorphism maintained by negative frequency dependent selection (NFDS) in a large ancestral population that becomes randomly fixed in bottlenecking encountered during budding speciation. B) Polymorphism maintained by NFDS that becomes fixed due to positive selection (PS) in a budding descendant that has dispersed into a new environment. Lineage widths in budding lineage diagrams represent effective population size. Under scenario A, variation is filtered randomly by drift into the descendant lineages. Under scenario B, the new regime rapidly fixes one allele/character state according to its new selective landscape.

**Table 1.**
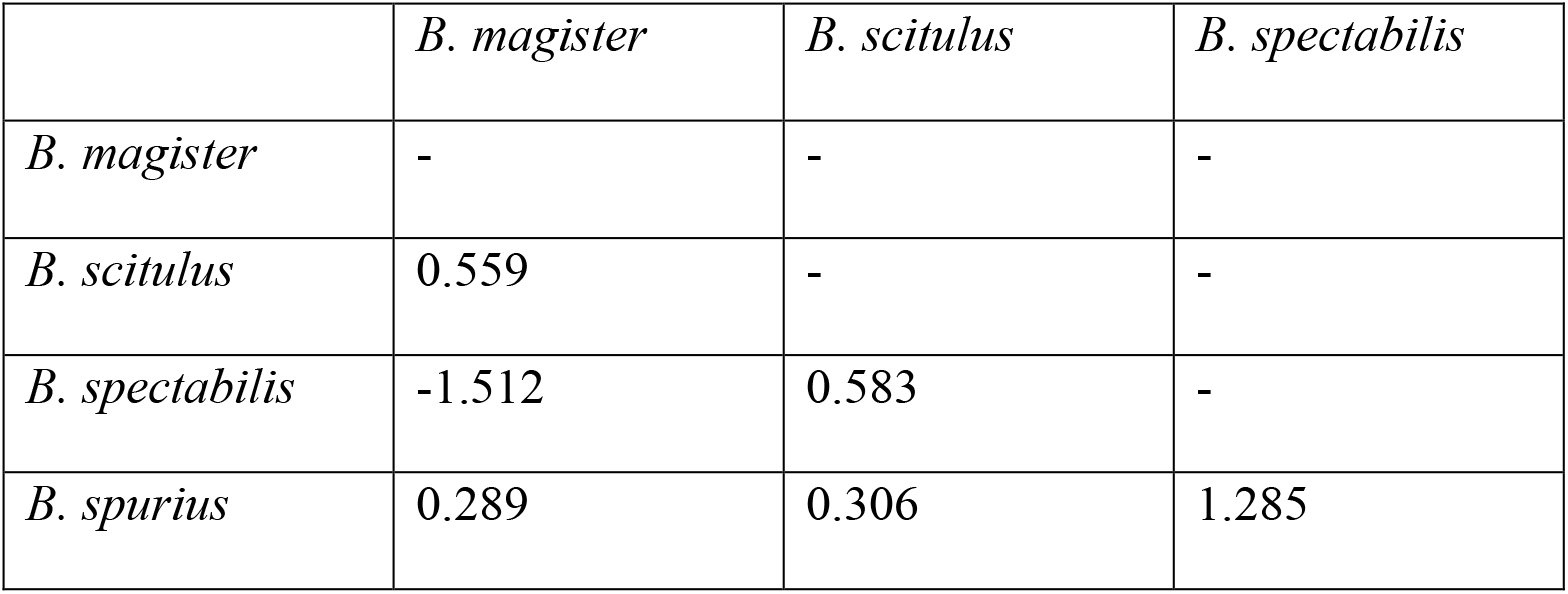
Pairwise phenotypic distances calculated among characters sorted from polymorphisms present in the ancestor, *Barycrinus rhombiferus* between descendant lineages, relative to null expectation generated under random sorting of ancestral polymorphisms. No pairs were statistically significant at the 2.5% threshold.

|  | *B. magister* | *B. scitulus* | *B. spectabilis* |
| --- | --- | --- | --- |
| *B. magister* | - | - | - |
| *B. scitulus* | 0.559 | - | - |
| *B. spectabilis* | -1.512 | 0.583 | - |
| *B. spurius* | 0.289 | 0.306 | 1.285 |

**Table 2.**
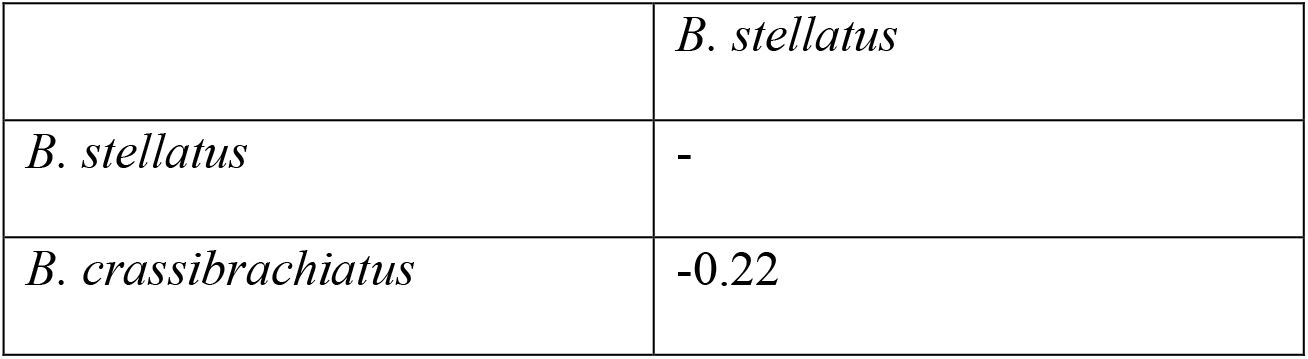
Pairwise phenotypic distances calculated among characters sorted from polymorphisms present in the ancestor, *Barycrinus spectabilis* between descendant lineages, relative to null expectation generated under random sorting of ancestral polymorphisms. No pairs were statistically significant at the 2.5% threshold.

Throughout the radiation of *Barycrinus*, variation maintained within ancestral lineages filtered down through a series of successively less variable descendants (Fig. 2). The only exceptions lie in the emergence of *B. rhombiferus* from *C. ibericus* (Kammer 2001) and *C. rex* (McIntosh 1984) from the hypothetical descendant of *C. ibericus*. The sudden and almost coincident emergence of high phenotypic variability in *B. rhombiferus* and *C. rex* from the monomorphic *C. ibericus* can be explained by a difference in sample size. *C. ibericus* is known only from a single specimen (Kammer 2001) and so the true extent of intraspecific variation that originated across this transition cannot be evaluated from the data used here. The descendants of polymorphic ancestors are universally less variable than their ancestors. In all, seven total lineages formed across *Barycrinus* by filtering variation maintained over millions of years by *B. rhombiferus* and *B. spurius*. While some of the variation displayed by *B. rhombiferus* may not have persisted over its entire range, the lineage did maintain high levels of polymorphism throughout its existence. Each descendant lineage that budded from *B. rhombiferus* and *B. spurius* might be characterized as a unique natural experiment, randomly fixing ancestral variation and either persisting or perishing in its new habitat. More ecological information would be needed to understand the specific phenotypic determinants of persistence and extinction among the varied *Barycrinus* offspring lineages.

### A brief note on speciation mode

The phylogenetic results in both *Ectocion* and *Barycrinus* reveal a pattern of extensive budding speciation. The relative frequency of alternative speciation modes in the fossil record is still not yet well known. Researchers have dealt with this uncertainty through a range of approaches. Some studies have explored patterns across a range of potential modes (Foote 1996), others have assumed budding *a priori* (Raup and Gould 1974, Van Valen 1975, Raup 1985), and still others have resorted to the use of cladograms, which make no assumptions about mode, at the cost of evolutionary specificity. Nevertheless, several studies have examined the frequency of modes across lineages. In general, anagenesis is thought to be rare relative to budding, occurring in a small minority of cases at one extreme (2%, Bapst and Hopkins 2017) and a larger minority at another (25%, Archibald 1993). Another important study found budding to form the predominant speciation pattern, with both anagenesis and bifurcating cladogenesis providing poor explanations for phylogenetic patterns in the fossil record (Wagner and Erwin 1995). Here, I assume that budding speciation forms the predominant pattern in both lineages analyzed, while also allowing for the potential of both bifurcating cladogenesis and anagenesis. The potential for anagenesis appears quite low across both datasets, with only only three candidate anagenetic ancestor-descendant species pairs: *E. osbornianus-E. superstes, B. stellatus-B. punctus, and B. rhombiferus-B. spectabilis*, due to their non-overlapping ranges. Each of these three species transitions could be explained by either anagenesis or budding. The results recovered here are robust to a small handful of anagenetic transitions, with only minor modifications of the interpretation of the demographic processes underlying ancestral (pseudo-) extinction and descendant speciation. The general result that ancestors tend to be more variable than descendants and that ancestral variation tends to sort randomly into descendants would also remain intact.

### Phenotypic variation, geographic range, and lineage duration

The pattern uncovered here of long lived, variable ancestors giving rise to multiple shorter lived, less variable descendants may be reflective of a more general phenomenon. *Barycrinus rhombiferus*, which gave rise to multiple descendant lineages over a long duration, was phenotypically variable, geographically widespread, and highly abundant (Gahn and Kammer 2002). The finding that descendant lineages displayed lower phenotypic variation is not surprising, given that they were also more geographically constrained and fewer in number than their ancestor. This is consistent with the demographic asymmetry associated with budding speciation and the peripheral geographic isolation expected under peripatric speciation (Mayr 1963). As a result, the pattern uncovered here highlights the potential role geography may play in shaping both patterns in lineage survivorship and the origin and maintenance of phenotypic variation across lineage radiations. However, while wide geographic range and high phenotypic variability have previously been linked to longer lineage durations (Liow 2007, Payne and Finnegan 2007), drawing causative links between these variables has proved challenging (Foote et al. 2008). It is therefore unclear whether *B. rhombiferus*’ lineage persistence can be explained by its high morphological variability and/or wide geographic range, or vice versa. A stronger understanding of the links between geographic range and morphological variability is also needed. While all three variables are highly conflated, in the clades examined here, incipient species form as subsets of the geographic range and stock of variation present in the ancestor. Future work will be needed to better explain the links between morphological variation, geographic range, and lineage duration in *Barycrinus* and other lineages.

### Budding, phylogenetic pattern, and paleobiological process

A pattern of multiple descendants arising from a single, long lived, ancestor was uncovered in both *Ectocion* and botyrocrinids. Such complex relationships cannot be represented meaningfully in a purely bifurcating cladogram, resulting only in the appearance of “rogue” taxa and polytomies. This shortcoming of using cladograms as a proxy of evolutionary relationships in paleobiology has long been recognized (e.g., Wagner and Erwin 1995, Bapst 2013). Biological reality demands the accommodation of diverse speciation modes when reconstructing phylogeny in the fossil record. Moving forward, it is important to consider how diverse biological processes such as multiple buddings, anagenesis, and even hybridization (Ausich and Meyer 1992) might shape relationships among fossil lineages. Although substantial advances have been made recently in the evaluation and representation of a greater diversity of speciation modes (Bapst and Hopkins 2017, Stadler et al. 2018, Parins-Fukuchi et al. 2019, Wright et al. 2021, Wright et al. 2022), many of these still operate from the premise that most relationships tend to be bifurcating and that hypothetical ancestors predominately link fossil lineages. This is largely a matter of implementation, stemming from the fact that most advances in phylogenetic tree searching over the past several decades have focused on inferring trees between contemporaneous lineages and perhaps also the pattern cladists’ denial of the identifiability of ancestral taxa (Engelmann and Wiley 1977). In the absence of overturning information (which may also exist) it is simply more parsimonious to initially assume hypotheses of direct ancestry, since these do not demand the *ad-hoc* construction of unobserved “ghost” lineages to explain phylogeny (Polly 1997). As a result, paleobiology might benefit from algorithms that begin from the premise that taxa originating at different times represent ancestor-descendent sequences *a priori* and allow character data to overturn this starting assumption when necessary.

Gahn and Kammer (2002) provided an excellent example of how a cladistic analysis might be supplemented with extensive natural history knowledge to derive deep evolutionary insights. While developing a thorough understanding of the full context and nuances of specimen-based data forms the foundation for sound paleobiological research, such insights can be supported by quantitative evidence and thorough graphical explorations. Once biological patterns can be reconstructed with a reasonably high degree of confidence, it becomes possible to narrow interpretations into a smaller range of possible explanatory, process-driven, scenarios (Fisher 1981). Evidence supporting alternative scenarios consistent with the reconstructed pattern might then be weighed using statistical criteria. Nevertheless, the first step is to further develop theory and methods supporting the reconstruction of ancestor-descendant relationships. While much important work has already been done in this area (see references in methods section), the results here especially highlight the importance of a continued rethinking of paleobiological phylogenetics and encourage a further untethering of the field from the shackles of cladistic dogma.

### Ancestral polymorphism, parallel speciation, and adaptive radiation

A major issue in the biology of adaptive radiation has been understanding how ecology shapes the phenotypes displayed by incipient species in the process and aftermath of adaptive radiation. Several authors have highlighted the role of parallel evolution in shaping these episodes (Schluter and Nagel 1995). Under this scenario, rapidly forming species evolve similar traits as they radiate into similar ecological niches. This model of adaptive radiation has been thought to be incompatible with the classic model of peripatric speciation (Mayr 1963), wherein species form rapidly during founder events. Paleobiologists might be inclined to liken these scenarios to budding speciation. The incompatibility between these two models presumably stems from the small population sizes displayed by the descendants that result from budding speciation, which would decrease the efficiency with which natural selection within each new population fixes similar variants in parallel among newly formed species.

While many examples of parallel speciation exist in the neontological literature, the paleobiological literature remains rife with examples of budding, suggesting a role for peripatry in the formation of new lineages. Careful examination of how variation is filtered and maintained at different taxonomic levels across varying ecological contexts might provide the key to reconciliation between these apparently conflicting views. Adaptation that occurs when populations move into new habitats can be fueled by large pools of standing ancestral variation (e.g., McGee 2020). If selection is strong enough among the descendant lineages, advantageous alleles may still overcome the effects of drift in small, peripherally isolated populations, allowing ancestral variation to fix in beneficial ways (Fig. 3b). This demands that the ancestral and descendant lineages exist in distinct ecological conditions, since the maintenance of variation in the ancestral population would require a different, balancing, selective regime than the positively-selective descendants. This type of dynamic may be shown between large, marine, ancestral populations of sticklebacks and their smaller descendant freshwater populations (e.g., Schluter and Conte 2009). Species selection also may be able to generate a similar pattern when the effect of drift is too great for selection to operate at the population level. Variation in peripherally-isolated populations might become fixed randomly, with only populations that stochastically fixed beneficial alleles able to persist. The *Barycrinus* analyses here are more consistent with the latter scenario, which showed that the pattern of character state sorting in descendant lineage cannot be distinguished from drift. Each of these alternatives would yield similar patterns to parallel speciation, with the only difference being the level at which variation filters out deleterious variation. While, in sticklebacks, population-level positive selection may have driven the appearance of parallel speciation, the *Barycrinus* analysis illustrates the possible role for species selection in generating similar patterns by differentially culling descendant lineages that each inherited a random suite of characters from their ancestor.

### Long term maintenance of polymorphisms

The trans-specific maintenance of polymorphic characters creates a natural link between the population processes maintaining phenotypic and genetic variation and the macroevolutionary processes shaping the evolution of lineages. Each of the datasets examined here displays polymorphisms maintained over long timescales, both within and between species. The polymorphisms present within lineages were confirmed or implied by the original studies to have been present throughout the duration of each lineage, even though trait frequencies may have oscillated over time in some cases. The true temporal persistence of these polymorphisms was also supported by their frequent appearance across species boundaries. While polymorphisms often sorted into monomorphic descendant lineages, they also persisted in several cases, such as the transition from *Ectocion mediotuber* to *Ectocion osbornianus* or *Bary-crinus rhombiferus* and *Barycrinus spurius*. Patterns in the maintenance and sorting of polymorphic traits across species boundaries in *Barycrinus* highlight how variation at the population level stochastically generates variation between species. The transformation of intrapopulation into trans-specific variation paves the way for higher-level processes, such as species selection (Stanley 1975), to operate. Because evolution is fundamentally rooted in the study of biological variation, it makes sense that understanding its persistence across scales provides the indispensable link between population processes and paleobiological patterns. Further developing this framework may provide the key to developing a true understanding of macroevolutionary *process* and how it connects to the mechanisms of population genetics.

### Linking population and macroevolutionary processes

When maintained over long periods against a consistent backdrop of balancing selection, lineages that maintain polymorphic variation will be better suited to persist compared to lineages that lose diversity to drift. It has been argued that the basic prerequisite for selection to operate above the species level is that phenotypic variation be fixed within lineages, but variable between lineages (Stearns 1986). However, the persistence of polymorphisms maintained over long periods within and distributed between fossil lineages adds additional complexity. When balancing selection is the maintaining force, even lineages that display variation may be subject to species selection under certain conditions. These polymorphic lineages display what might be viewed as a lineage-level analogue to heterozygote advantage (left side of Fig. 4). Under this scenario, bet-hedging in the face of spatial or temporal variation in environment and/or NFDS at the population level maintains stability in trait or genotype frequencies that facilitates higher-level dynamics, such as species selection, to operate simultaneously. In this case, the necessary condition for species selection to operate would be ecologically-induced maintenance of polymorphism (potentially both within and across lineages), rather than a total lack of intraspecific variation. On the other hand, lineages that fix variation from polymorphic ancestors may achieve greater persistence by escaping into a new ecological niche that does not impose balancing selection (Fig. 4). Overall, it is important to consider both inter- and intraspecific variation and the ecological contexts within which they are maintained and sorted among lineages when exploring higher-level evolutionary dynamics.

**Figure 4.**
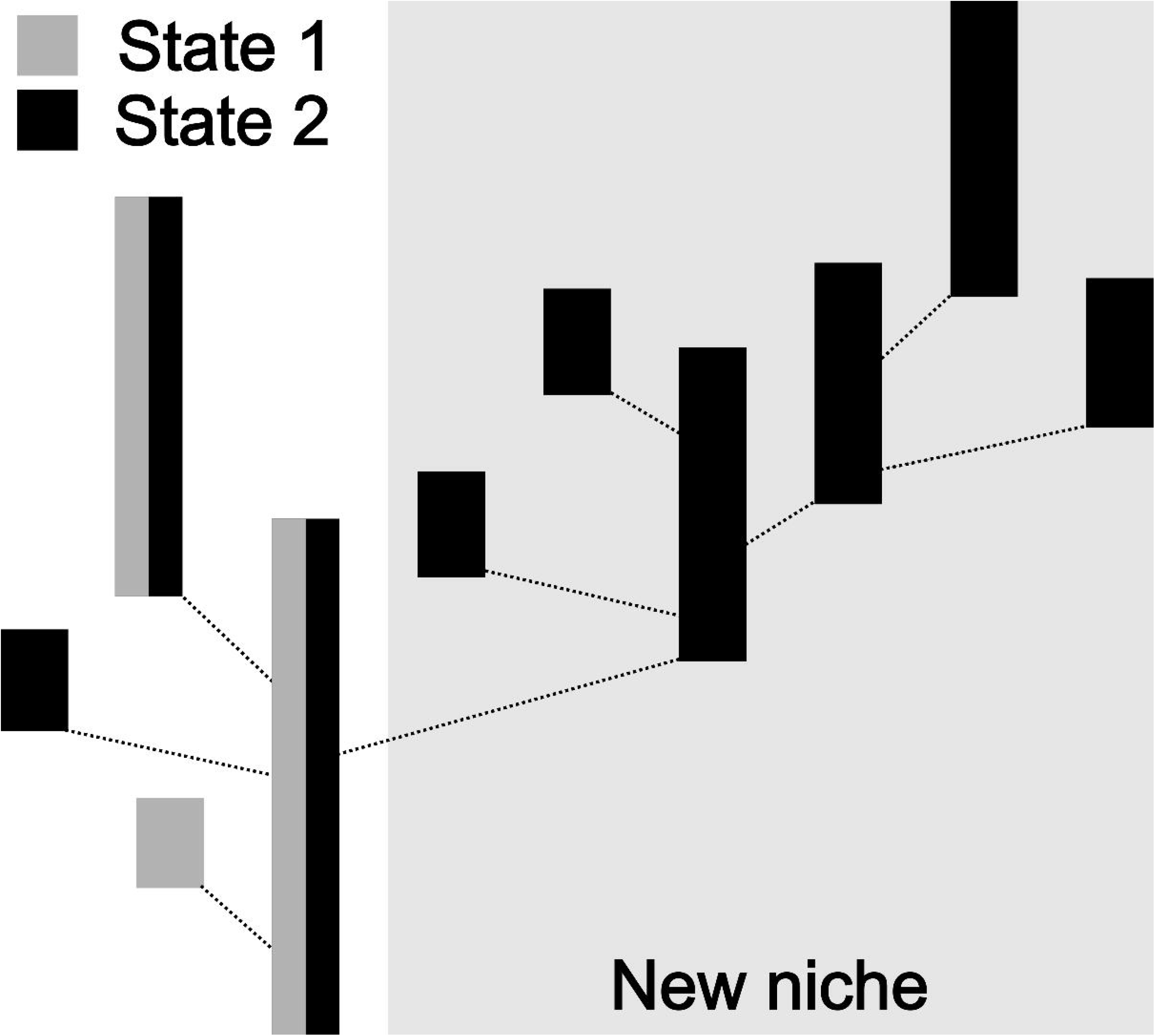
Species selection dynamics stemming from a persistently polymorphic ancestral population. In this hypothetical scenario, negative frequency-dependent selection forms the selective background in the ancestor. When descendant lineages randomly fix this ancestral variation as they originate, they demonstrate low survivorship in the ancestral niche. Only lineages that can escape and radiate into a new niche while fixing ancestral variation display high survivorship. Ecological opportunity afforded by the new niche may even facilitate enhanced lineage survivorship and proliferation if the trait fixed in the descendant is congruent with the selective demands of the new habitat.

While the level at which variation is displayed remains a critical point in identifying the boundary conditions under which higher-level selection can operate, perhaps an even more crucial point lies in considering the selective and ecological contexts occupied by different lineages. Rather than demanding the absence of variation within species, the conditions allowing species-level selection may instead be specified by comparing the distribution of variation within species (polyvs. monomorphic) to the ecological conditions occupied across species (e.g., does competition or predation universally impose frequency-dependent fitness effects within species?). The longer durations exhibited by *B. rhombiferus* and *B. spurius* may stem from higher species-level fitness for polymorphic taxa, which could be driven by balancing forces such as environmental variability or negative frequency-dependence resulting from competition or predation. If intra-specific polymorphism is maintained by environmental heterogeneity associated with more generalist ecologies, the new niches occupied by successful monomorphic lineages should tend to be more specialized. This would yield distinct macroevolutionary dynamics, with generalist polymorphic lineages displaying high lineage survivorship and specialist monomorphic lineages displaying rapid species turnover. This dynamic would explain the persistence of polymorphism in the fossil record by the ecological dynamics displayed by Mississippian crinoids (Kammer et al. 1997). Nevertheless, moving forward, it will also be important to test such hypotheses within the context of geographic range and abundance, which may also impact lineage duration and turnover patterns (Liow 2007).

It has long been hypothesized that biased patterns of extinction among new species might shape macroevolutionary patterns observed in the fossil record by culling “ephemeral” species before they are able to fossilize (Raup and Stanley 1978; pp. 105, Stanley 1979, Rosenblum et al. 2012, Rabosky 2013). Such biased extinction patterns could arise when new lineages are unable to adapt rapidly enough to new environmental conditions (De Lisle et al. 2021). Meanwhile, paleontological work has suggested that high morphological variation within species might help fuel lineage radiations (Webster 2007). If drawn from highly-diverse ancestral populations, the variation displayed by rapidly radiating descendant lineages will become sorted through a mix of stochasticity and adaptation, potentially occurring at multiple levels. Incipient species that are able to draw upon a large well of ancestral variation may stand a greater chance of inheriting “rescue” alleles that enable them to escape extinction long enough to avoid non-preservation in the fossil record due to ephemerality.

The “natural experiments” displayed by descendant *Barycrinus* species illustrate how filtering of ancestral variation, a process that occurs at the population level, might form the basic conditions and raw material for macroevolutionary processes, such as species selection, to operate. Random sorting of ancestral variation into descendant lineages is a pattern predicted by classic evolutionary theory (e.g., Simpson 1944, Mayr 1963, Wright 1982). The manner by which this occurs, when compared against the range of selective conditions inhabited by incipient lineages, might explain how variation at the population level in highly-diverse ancestral populations might filter into descendant lineages to shape macroevolutionary dynamics. Recent work has suggested that gene tree discordance, which is at least partially caused by incomplete lineage sorting (ILS)-- the random sorting of ancestral variation into descendant lineages--is greater during periods of rapid phenotypic innovation, such as during the early evolution of mammals, birds, or angiosperms (Parins-Fukuchi et al. 2021). This link implies high genetic diversity (and thus perhaps high phenotypic variation as well) within lineages during the early stages of rapid clade diversification. Such variation will stand a good chance of sorting in ways that are discordant with the order of species divergences, yielding two main effects: 1) high gene-tree discordance and 2) the apparent appearance of rapid morphological variation distributed haphazardly across lineages. The resulting combination of stochastic sorting, parallel speciation, and species selection may yield the extensive gene-tree discordance and hemiplasy (Avise 2008, Gurrero and Hahn 2018) that often accompanies rapid radiations. Future work drawing more explicit links between coalescent expectations under ILS and the sorting of phenotypic ancestral variation across fossil lineages may help shed light on how population processes provide the fuel for large-scale macroevolutionary processes and patterns to operate.

### Conclusion

Explaining the patterns observed by paleontologists in the fossil record in terms of population processes has been a major goal of evolutionary paleobiology since the modern synthesis (e.g., Simpson 1944, Kermack 1954, Van Valen 1963, Eldredge and Gould 1972, Fisher 1985). The persistence of phenotypic variation displayed across fossil lineages, while interesting from a population genetic perspective, may provide even more groundbreaking insights when considered from a macroevolutionary perspective. The patterns reconstructed here highlight how further advances in how we model 1) speciation dynamics, including ancestor-descendant relationships and 2) the filtering of polymorphic phenotypic variation can vastly increase the scope of evolutionary questions we are able to evaluate in the fossil record. More work will also be needed at the population level to better understand how frequencies in polymorphic traits evolve across the stratigraphic ranges of continuous populations. This will contribute to a stronger quantitative understanding of the processes that maintain biological variation among long-lived fossil lineages, such as *Ectocion osbornianus* or *Barycrinus rhombiferus*. Scaling up, understanding how pools of maintained phenotypic variation segregate during between incipient species can provide a crucial link between the mechanisms explored by population genetics and the patterns historically explored in evolutionary paleobiology. Developing stronger links across these levels can help provide more cohesive and deeper explanations of how population-level evolutionary change scales to the luxuriant diversity of patterns observed throughout the history of life.

## Acknowledgements

Several of the core themes present in this manuscript were sparked by a series of engaging conversations with L. Rowe. J. Saulsbury offered much salient guidance on the interpretation of the evolutionary morphology and stratigraphy of fossil crinoids as well as excellent discussion around the macroevolutionary points of the manuscript. N. Walker-Hale offered helpful criticism of the manuscript. I thank M. Ahmad-Gawel for being a constant sounding board and critical discussion partner for my more audacious hypotheses of evolutionary process and for offering her keen and skilled eye for editing. M. Foote provided insightful criticism of the manuscript and ideas presented herein. The manuscript was greatly improved by reviews from F. Gahn and D. Wright, who each provided extensive and generous insights into cladid crinoid paleobiology and evolutionary processes. Any errors or overwrought biological interpretations that remain lie solely on my own shoulders. I was supported by National Science Foundation grant DEB-2217117 throughout the course of this work.

## References

Alroy, J., 1995. Continuous track analysis: a new phylogenetic and biogeographic method. Systematic Biology, 44(2), pp.152–178.

Antonovics, J. and Kareiva, P., 1988. Frequency-dependent selection and competition: empirical approaches. Philosophical Transactions of the Royal Society of London. B, Biological Sciences, 319(1196), pp.601–613.

Archibald, J.D., 1993. The importance of phylogenetic analysis for the assessment of species turnover: a case history of Paleocene mammals in North America. Paleobiology, 19(1), pp.1–27.

Ausich, W.I., 1980. A model for niche differentiation in Lower Mississippian crinoid communities. Journal of Paleontology, pp.273–288.

Ausich, W.I. and Bottjer, D.J., 1982. Tiering in suspension-feeding communities on soft substrata throughout the Phanerozoic. Science, 216(4542), pp.173–174.

Ausich, W.I. and Meyer, D.L., 1994. Hybrid crinoids in the fossil record (Early Mississippian, Phylum Echinodermata). Paleobiology, 20(3), pp.362–367.

Avise, J.C. and Robinson, T.J., 2008. Hemiplasy: a new term in the lexicon of phylogenetics. Systematic biology, 57(3), pp.503–507.

Aze, T., Ezard, T.H., Purvis, A., Coxall, H.K., Stewart, D.R., Wade, B.S. and Pearson, P.N., 2011. A phylogeny of Cenozoic macroperforate planktonic foraminifera from fossil data. Biological Reviews, 86(4), pp.900–927.

Bapst, D.W., 2013. When can clades be potentially resolved with morphology?. PLoS One, 8(4), p.e62312.

Bapst, D.W. and Hopkins, M.J., 2017. Comparing cal3 and other a posteriori time-scaling approaches in a case study with the pterocephaliid trilobites. Paleobiology, 43(1), pp.49–67.

Barrett, R.D. and Schluter, D., 2008. Adaptation from standing genetic variation. Trends in ecology & evolution, 23(1), pp.38–44.

Benton, M.J. and Pearson, P.N., 2001. Speciation in the fossil record. Trends in Ecology & Evolution, 16(7), pp.405–411.

Brothwell, D.R. ed., 2014. Dental Anthropology: Volume V: Society for the Study of Human Biology. 5, pp.286. Elsevier.

Chakraborty, M. and Fry, J.D., 2016. Evidence that environmental heterogeneity maintains a detoxifying enzyme polymorphism in Drosophila melanogaster. Current Biology, 26(2), pp.219–223.

Charlesworth, D., 2006. Balancing selection and its effects on sequences in nearby genome regions. PLoS Genetics, 2(4), p.e64.

Christiansen, F.B., 1975. Hard and soft selection in a subdivided population. The American Naturalist, 109(965), pp.11–16.

Clarke, B.C., 1979. The evolution of genetic diversity. Proceedings of the Royal Society of London. Series B. Biological Sciences, 205(1161), pp.453–474.

Cole, S.R., 2021. Hierarchical controls on extinction selectivity across the diplobathrid crinoid phylogeny. Paleobiology, 47(2), pp.251–270.

Cole, S.R., Wright, D.F. and Ausich, W.I., 2019. Phylogenetic community paleoecology of one of the earliest complex crinoid faunas (Brechin Lagerstätte, Ordovician). Palaeogeography, Palaeoclimatology, Palaeoecology, 521, pp.82–98.

Cope, E.D., 1882. Contributions to the history of the Vertebrata of the Lower Eocene of Wyoming and New Mexico, made during 1881. Proceedings of the American Philosophical Society, 20(111), pp.139–197.

De Lisle, S.P., Punzalan, D., Rollinson, N. and Rowe, L., 2021. Extinction and the temporal distribution of macroevolutionary bursts. Journal of Evolutionary Biology, 34(2), pp.380–390.

Dijkstra, P.D. and Border, S.E., 2018. How does male–male competition generate negative frequency-dependent selection and disruptive selection during speciation?. Current Zoology, 64(1), pp.89–99.

Engelmann, G.F. and Wiley, E.O., 1977. The place of ancestor-descendant relationships in phylogeny reconstruction. Systematic Biology, 26(1), pp.1–11.

Eldredge, N. and Gould, S.J., 1972. Punctuated equilibria: an alternative to phyletic gradualism. Models in Paleobiology, 1972, pp.82–115.

Fisher, D.C., 1991. Phylogenetic analysis and its application in evolutionary paleobiology. Short Courses in Paleontology, 4, pp.103–122.

Fisher, D.C., 2008. Stratocladistics: integrating temporal data and character data in phylogenetic inference. Annual Review of Ecology, Evolution, and Systematics, 39, pp.365–385.

Fitzpatrick, M.J., Feder, E., Rowe, L. and Sokolowski, M.B., 2007. Maintaining a behaviour polymorphism by frequency-dependent selection on a single gene. Nature, 447(7141), pp.210–212.

Fisher, D.C., 1981. The role of functional analysis in phylogenetic inference: examples from the history of the Xiphosura. American Zoologist, 21(1), pp.47–62.

Fisher, D.C., 1985. Evolutionary morphology: beyond the analogous, the anecdotal, and the ad hoc. Paleobiology, 11(1), pp.120–138.

Foote, M., 1996. On the probability of ancestors in the fossil record. Paleobiology, 22(2), pp.141–151.

Foote, M., Crampton, J.S., Beu, A.G. and Cooper, R.A., 2008. On the bidirectional relationship between geographic range and taxonomic duration. Paleobiology, 34(4), pp.421–433.

Gallet, R., Froissart, R. and Ravigné, V., 2018. Experimental demonstration of the impact of hard and soft selection regimes on polymorphism maintenance in spatially heterogeneous environments. Evolution, 72(8), pp.1677–1688.

Gahn, F.J. and Kammer, T.W., 2002. The cladid crinoid Barycrinus from the Burlington Limestone (early Osagean) and the phylogenetics of Mississippian botryocrinids. Journal of Paleontology, 76(1), pp.123–133.

Gahn, F.J., 2003. A stratocladistic approach for evaluating hypotheses of cladogenesis, anagenesis, and budding applied to the Mississippian crinoid *Barycrinus*. Geological Society of America annual meeting. Seattle, WA, USA. 35(6), pp. 166.

Gilbert, C.C., 2013. Cladistic analysis of extant and fossil African papionins using craniodental data. Journal of human evolution, 64(5), pp.399–433.

Gingerich, P.D., 1979. The stratophenetic approach to phylogeny reconstruction in vertebrate paleontology. In Phylogenetic analysis and paleontology (pp. 41–78). Columbia University Press. New York.

Gingerich, P.D., 1974. Stratigraphic record of early Eocene Hyopsodus and the geometry of mammalian phylogeny. Nature, 248(5444), pp.107–109.

Goldberg, E.E., Kohn, J.R., Lande, R., Robertson, K.A., Smith, S.A. and Igić, B., 2010. Species selection maintains self-incompatibility. Science, 330(6003), pp.493–495.

Granger, W. 1915. Part III.--Order Condylarthra. Families Phenacodontidae and Meniscotheriidae, p. 329–361. In W. D. Matthew and W. Granger (eds.), A revision of the lower Eocene Wasatch and Wind River Faunas. Bulletin of the American Museum of Natural History, 39.

Guerrero, R.F. and Hahn, M.W., 2017. Speciation as a sieve for ancestral polymorphism. Molecular Ecology, 26(20), pp.5362–5368.

Guerrero, R.F. and Hahn, M.W., 2018. Quantifying the risk of hemiplasy in phylogenetic inference. Proceedings of the National Academy of Sciences, 115(50), pp.12787–12792.

Hall, J. 1858. Report on the Geological Survey of Iowa embracing the results of investigations made during portions of the years 1855, 1856, 1857. Geological Survey of Iowa, 1(1-2), pp. 724.

Igic, B., Bohs, L. and Kohn, J.R., 2006. Ancient polymorphism reveals unidirectional breeding system shifts. Proceedings of the National Academy of Sciences, 103(5), pp.1359–1363.

Jackson, J.B. and Cheetham, A.H., 1994. Phylogeny reconstruction and the tempo of speciation in cheilostome Bryozoa. Paleobiology, 20(4), pp.407–423.

Kammer, T.W., 2001. Phenotypic bradytely in the Costalocrinus-Barycrinus lineage of Paleozoic cladid crinoids. Journal of Paleontology, 75(2), pp.383–389.

Kammer, T.W., Baumiller, T.K. and Ausich, W.I., 1997. Species longevity as a function of niche breadth: evidence from fossil crinoids. Geology, 25(3), pp.219–222.

Kermack, K.A., 1954. A biometrical study of Micraster coranguinum and M.(Isomicraster) senonensis. Philosophical Transactions of the Royal Society of London. Series B, Biological Sciences, pp.375–428.

Lai, Y.T., Yeung, C.K., Omland, K.E., Pang, E.L., Hao, Y.U., Liao, B.Y., Cao, H.F., Zhang, B.W., Yeh, C.F., Hung, C.M. and Hung, H.Y., 2019. Standing genetic variation as the predominant source for adaptation of a songbird. Proceedings of the National Academy of Sciences, 116(6), pp.2152–2157.

Liow, L.H., 2007. Does versatility as measured by geographic range, bathymetric range and morphological variability contribute to taxon longevity?. Global Ecology and Biogeography, 16(1), pp.117–128.

Mayr, E. 1963. Animal species and evolution. Harvard University Press, Cambridge.

McDonald, J.F. and Ayala, F.J., 1974. Genetic response to environmental heterogeneity. Nature, 250(5467), pp.572–574.

McGee, M.D., Borstein, S.R., Meier, J.I., Marques, D.A., Mwaiko, S., Taabu, A., Kishe, M.A., O’Meara, B., Bruggmann, R., Excoffier, L. and Seehausen, O., 2020. The ecological and genomic basis of explosive adaptive radiation. Nature, 586(7827), pp.75–79.

McIntosh, G.C., 1984. Devonian cladid inadunate crinoids: family Botryocrinidae Bather, 1899. Journal of Paleontology, pp.1260–1281.

Meek, F.B., and Worthen, A.H. 1860. Description of new species of Crinoidea and Echinoidea from the Carboniferous rocks of Illinois, and other western states. Academy of Natural Sciences, Philadelphia, Proceedings, Series 2(4), pp.379–397.

Meek, F.B. and Worthen, A.H., 1870. Descriptions of new species and genera of fossils from the Palaeozoic rocks of the western states. Proceedings of the Academy of Natural Sciences of Philadelphia, pp.22–56.

Owen, D.D. and Shumard, B.F., 1852. Descriptions of seven new species of crinoidea from the subcarboniferous of Iowa and Illinois. Journal of the Academy of Natural Sciences of Philadelphia, series, 2(2), pp.89–94.

Parins-Fukuchi, C.T., Stull, G.W. and Smith, S.A., 2021. Phylogenomic conflict coincides with rapid morphological innovation. Proceedings of the National Academy of Sciences, 118(19), p.e2023058118.

Parins-Fukuchi, C.T., 2021. Morphological and phylogeographic evidence for budding speciation: an example in hominins. Biology Letters, 17(1), p.20200754.

Parins-Fukuchi, C., Greiner, E., MacLatchy, L.M. and Fisher, D.C., 2019. Phylogeny, ancestors, and anagenesis in the hominin fossil record. Paleobiology, 45(2), pp.378–393.

Payne, J.L. and Finnegan, S., 2007. The effect of geographic range on extinction risk during background and mass extinction. Proceedings of the National Academy of Sciences, 104(25), pp.10506–10511.

Pease, J.B., Haak, D.C., Hahn, M.W. and Moyle, L.C., 2016. Phylogenomics reveals three sources of adaptive variation during a rapid radiation. PLoS biology, 14(2), p.e1002379.

Polly, P.D., 1997. Ancestry and species definition in paleontology: a stratocladistic analysis of Paleocene-Eocene Viverravidae (Mammalia, Carnivora) from Wyoming. Contributions from the Museum of Paleontology. 30(1), pp.1–53.

Prout T., 2000. How well does opposing selection maintain variation? In: Singh RS, Krimbas CB, editors. Evolutionary genetics: from molecules to morphology. Cambridge: Cambridge University Press. pp.157–181.

Rabosky, D.L., 2013. Diversity-dependence, ecological speciation, and the role of competition in macroevolution. Annual Review of Ecology, Evolution, and Systematics, 44, pp.481–502.

Raup, D.M. and Gould, S.J., 1974. Stochastic simulation and evolution of morphology-towards a nomothetic paleontology. Systematic Biology, 23(3),pp.305–322.

Raup, D. and Stanley, S.M., 1978. Principles of Paleontology. Macmillan.

Raup, D.M., 1985. Mathematical models of cladogenesis. Paleobiology, 11(1), pp.42–52.

Rensch, B., 1959. 6. The Rules of Kladogenesis (Phylogenetic Branching). In Evolution Above the Species Level (pp. 97-280). Columbia University Press.

Rosenblum, E.B., Sarver, B.A., Brown, J.W., Des Roches, S., Hardwick, K.M., Hether, T.D., Eastman, J.M., Pennell, M.W. and Harmon, L.J., 2012. Goldilocks meets Santa Rosalia: an ephemeral speciation model explains patterns of diversification across time scales. Evolutionary Biology, 39, pp.255–261.

Russell, L.S., 1929. Paleocene vertebrates from Alberta. American Journal of Science, 5(98), pp.162–178.

Saitou, N. and Nei, M., 1987. The neighbor-joining method: a new method for reconstructing phylogenetic trees. Molecular biology and evolution, 4(4), pp.406–425.

Schluter, D., 2000. Ecological character displacement in adaptive radiation. The American Naturalist, 156(S4), pp.S4–S16.

Schluter, D. and Conte, G.L., 2009. Genetics and ecological speciation. Proceedings of the National Academy of Sciences, 106(supp. 1), pp.9955–9962.

Schluter, D. and Nagel, L.M., 1995. Parallel speciation by natural selection. The American Naturalist, 146(2), pp.292–301.

Schopf, T.J. and Gooch, J.L., 1972. A natural experiment to test the hypothesis that loss of genetic variability was responsible for mass extinctions of the fossil record. The Journal of Geology, 80(4), pp.481–483.

Sicard, A., Kappel, C., Lee, Y.W., Woźniak, N.J., Marona, C., Stinchcombe, J.R., Wright, S.I. and Lenhard, M., 2016. Standing genetic variation in a tissue-specific enhancer underlies selfing-syndrome evolution in Capsella. Proceedings of the National Academy of Sciences, 113(48), pp.13911–13916.

Simpson, G.G., 1944. Tempo and Mode in Evolution. Columbia University Press. New York.

Stadler, T., Gavryushkina, A., Warnock, R.C., Drummond, A.J. and Heath, T.A., 2018. The fossilized birth-death model for the analysis of stratigraphic range data under different speciation modes. Journal of Theoretical Biology, 447, pp.41–55.

Stanley, S.M., 1975. A theory of evolution above the species level. Proceedings of the National Academy of Sciences, 72(2), pp.646–650.

Stanley SM. 1979. Macroevolution: Pattern and Process. San Francisco, CA: Freeman

Stearns, S.C., 1986. Natural selection and fitness, adaptation and constraint. In: Raup D. M. and Jablonski, D., eds., Patterns and Processes in the History of Life: Report of the Dahlem Workshop on Patterns and Processes in the History of Life, pp. 23–44. Springer Berlin Heidelberg.

Takahashi, Y., Yoshimura, J., Morita, S. and Watanabe, M., 2010. Negative frequency-dependent selection in female color polymorphism of a damselfly. Evolution, 64(12), pp.3620–3628.

Thewissen, J.G.M., 1990. Evolution of Paleocene and Eocene Phenacodontidae (Mammalia, Condylarthra). University of Michigan, Papers on Paleontology, 29, pp.1–10.

Thewissen, J.G.M., 1992. Temporal data in phylogenetic systematics: an example from the mammalian fossil record. Journal of Paleontology, 66(1), pp.1–8.

Thompson, K.A., Osmond, M.M. and Schluter, D., 2019. Parallel genetic evolution and speciation from standing variation. Evolution Letters, 3(2), pp.129–141.

Trueman, J.W., 2010. A new cladistic analysis of Homo floresiensis. Journal of human evolution, 59(2), pp.223–226.

Van Valen, L., 1969. Variation genetics of extinct animals. The American Naturalist, 103(931), pp.193–224.

Van Valen, L., 1975. Group selection, sex, and fossils. Evolution, pp.87–94.

Van Valen, L., 1963. Selection in natural populations: *Merychippus primus*, a fossil horse. Nature, 197, pp.1181–1183.

Wagner, P.J., 1998. A likelihood approach for evaluating estimates of phylogenetic relationships among fossil taxa. Paleobiology, 24(4), pp.430–449.

Wagner, P.J., 2000. Phylogenetic analyses and the fossil record: tests and inferences, hypotheses and models. Paleobiology, 26(S4), pp.341–371.

Wagner, P.J., Erwin, D.H., 1995. Phylogenetic patterns as tests of speciation models. In: Anstey, R.L, editor. New approaches to speciation in the fossil record. Columbia University Press, New York, pp.87–122.

Warnock, R.C., Heath, T.A. and Stadler, T., 2020. Assessing the impact of incomplete species sampling on estimates of speciation and extinction rates. Paleobiology, 46(2), pp.137–157.

Wiens, J.J., 1995. Polymorphic characters in phylogenetic systematics. Systematic Biology, 44(4), pp.482–500.

Wiens, J.J., 1999. Polymorphism in systematics and comparative biology. Annual Review of Ecology and Systematics, 30(1), pp.327–362.

Whiting, E.T., Steadman, D.W. and Vliet, K.A., 2016. Cranial polymorphism and systematics of Miocene and living Alligator in North America. Journal of Herpetology, 50(2), pp.306–315.

Webster, M., 2007. A Cambrian peak in morphological variation within trilobite species. Science, 317(5837), pp.499–502.

Wood, A.R., Zelditch, M.L., Rountrey, A.N., Eiting, T.P., Sheets, H.D. and Gingerich, P.D., 2007. Multivariate stasis in the dental morphology of the Paleocene-Eocene condylarth Ectocion. Paleobiology, 33(2), pp.248–260.

Wright, A.M., Bapst, D.W., Barido-Sottani, J. and Warnock, R.C., 2022. Integrating fossil observations into phylogenetics using the fossilized birth–death model. Annual Review of Ecology, Evolution, and Systematics, 53, pp.251–273.

Wright, A.M., Wagner, P.J. and Wright, D.F., 2021. Testing character evolution models in phylogenetic paleobiology: a case study with Cambrian echinoderms. Cambridge University Press. Cambridge.

Wright, D.F., 2022. Integrating probabilistic phylogenetics with the theory of punctuated equilibria to test the tempo and mode of speciation in the fossil record. Geological Society of America annual meeting. Denver, CO. Vol 54, No. 5, doi: 10.1130/abs/2022AM-380438

